# Why Ambrosia Beetles Keep Only One Fungus: Competitive Asymmetry, Bottleneck Drift, and Eco-Evolutionary Fixation

**DOI:** 10.64898/2025.12.18.694882

**Authors:** Zi-Ru Jiang

## Abstract

Ambrosia beetles rely on vertically transmitted fungal cultivars for nutrition, yet empirical studies reveal a persistent paradox: early galleries often host multiple fungal associates, whereas mature systems exhibit strict one-beetle–one-fungus specificity. The mechanisms driving this transition remain unresolved. Here we develop an integrative eco-evolutionary framework combining deterministic competition (Lotka–Volterra dynamics), stochastic drift, and severe mycangial transmission bottlenecks to quantify the stability and turnover of fungal symbionts. Across >50,000 simulation replicates, we show that multi-fungal coexistence is inherently transient under biologically realistic conditions. Even minimal competitive asymmetries (<5%) yield deterministic exclusion; repeated bottlenecks amplify drift and accelerate diversity loss; and weak but persistent selection ensures long-term fixation of the superior symbiont over hundreds of generations. Replacement by novel fungi follows a strict invasion threshold (s > 1/(2N)), consistent with classical fixation theory and matching empirical patterns of historical symbiont turnover. Multispecies communities collapse even more rapidly, persisting only over short ecological timescales. Together, these results provide a mechanistic explanation for the pervasive one-fungus specificity observed across ambrosia beetles. By unifying ecological competition, demographic stochasticity, and evolutionary stability theory, this work offers a general framework for understanding partner fidelity, symbiont filtering, and lineage turnover in insect–microbe mutualisms.

## Introduction

Ambrosia beetles (Curculionidae: Scolytinae and Platypodinae) maintain one of the most specialized and evolutionarily persistent nutritional mutualisms known among insects, relying entirely on cultivated fungi for larval and adult development. Classical foundational work first described mycangia—specialized organs enabling precise transport of fungal propagules—and documented remarkably strict fidelity between beetles and their associated fungi (Batra 1963, 1966; Francke-Grosmann 1956, 1967). Subsequent ecological and evolutionary syntheses further established that ambrosia beetles practice a unique form of “insect agriculture,” involving obligate fungal cultivation, anatomically structured transmission routes, and complete dependence on fungal metabolism for nutrition (Batra 1985; Mueller et al. 2001; Hulcr & Stelinski 2017; Jiang et al. 2021, 2022, 2023). More recent microbiome studies further reveal that ambrosia beetle symbioses include not only nutritional fungi but also diverse bacterial partners that may contribute to metabolism, defense, and community stability (e.g., Cambronero-Heinrichs et al. 2025). Experimental work also suggests that growth performance of ambrosia fungi can be strongly influenced by host tree species, introducing an environmental dimension to symbiosis persistence (Decker et al. 2025).

Despite this deep evolutionary history, modern molecular surveys reveal a striking combination of fidelity and flexibility. Community sequencing shows that although many beetle species maintain a single dominant fungal symbiont, early gallery phases often contain multiple co-occurring fungal associates (Kajimura & Hijii 1992; Kostovcik et al. 2015; Freeman et al. 2016; Jiang et al. 2021, 2022, 2023). These mixed communities undergo predictable successional transitions, with a single primary symbiont nearly always dominating by the time new adults emerge (Biedermann et al. 2013; Mayers et al. 2022). Anatomical specialization of mycangia likely contributes to this filtering: these structures restrict physical space, impose strong physiological selection, and favor only fungi with traits suited for transport and persistence (Mayers et al. 2015; Hulcr & Stelinski 2017).

At macroevolutionary scales, phylogenetic and genomic studies reveal that ambrosia symbioses are neither static nor immutable. Multiple fungal lineages show evidence of recurrent host switching, symbiont replacement, and adaptive divergence (Alamouti et al. 2009; Dreaden et al. 2014; Carrillo et al. 2014; O’Donnell et al. 2015). Genome-level analyses further demonstrate that ambrosia fungi undergo extensive metabolic specialization and gene-content remodeling, consistent with long-term coevolutionary filtering and adaptation to the beetle-associated niche (Blaz et al. 2018). Such findings indicate a dynamic evolutionary landscape in which functional specialization promotes fidelity, while ecological or genetic shifts occasionally open pathways for symbiont turnover. Recent genomic analyses further reveal pronounced diversification and lineage-specific remodeling of ambrosia fungi, consistent with repeated adaptive shifts within beetle-associated niches (Huang et al. 2025).

Yet, despite clear evidence of early-stage flexibility and long-term evolutionary change, most ambrosia beetle species maintain strict one-fungus specificity across generations (Hulcr & Dunn 2011; Six 2012). Current empirical knowledge does not fully explain why: Why are multi-fungal communities rare and short-lived? Why do nearly all systems converge on a single symbiont? Under what conditions can a novel fungus invade and replace a resident lineage?

Ecological studies of wood-decay fungi demonstrate that competitive hierarchies, interference competition, and priority effects strongly constrain coexistence (Boddy 2001; Hiscox et al. 2015). These mechanisms are expected to be even stronger within the spatially confined galleries of ambrosia beetles. Simultaneously, mycangial transmission imposes extreme population bottlenecks—often transferring only tens of propagules (Harrington & Fraedrich 2010)—which strongly amplify genetic drift. Work on microbial and endosymbiotic systems shows that repeated bottlenecks rapidly erode diversity, drive stochastic fixation, and enforce symbiont filtering (Moran & Sloan 2015; Wernegreen 2015). Parallel theoretical models of microbiome dynamics further demonstrate that interactions between drift and selection reliably collapse multi-species communities into monodominance (Chen et al. 2024).

Theoretical studies of mutualism evolution similarly predict that multi-partner associations are inherently unstable. Even slight differences in partner performance can be magnified by drift and selection, ultimately resolving into one-to-one associations (Foster & Kokko 2006). These predictions align with ambrosia systems but lack a unified quantitative framework linking competition, bottlenecks, stochastic drift, and evolutionary replacement.

Taken together, empirical evidence points toward a system shaped by ecological competitive filtering, strong demographic constraints, and evolutionary turnover across extended timescales. However, no existing work integrates these processes into a mechanistic framework capable of explaining: (1) why multi-fungal coexistence is transient, (2) how weak competitive differences still lead to deterministic long-term fixation, (3) under what conditions novel fungi can successfully invade established symbioses, (4) why single-symbiont dominance is nearly universal over evolutionary timescales.

To address these gaps, we develop an integrative ecological–evolutionary modeling framework that combines (i) competitive growth dynamics, (ii) stochastic drift under biologically realistic mycangial bottlenecks, and (iii) replicator-based stability analysis of evolutionary equilibria. Using this unified approach, we demonstrate that single-symbiont dominance emerges naturally from fundamental processes, providing the first mechanistic explanation for the pervasive one-beetle–one-fungus pattern across ambrosia beetles while also accounting for short-term coexistence and rare symbiont turnover events.

## METHODS

### Overview of the modeling framework

To investigate the evolutionary stability of fungal symbioses in ambrosia beetles, we developed an integrated simulation framework that combines (i) ecological competition among fungal symbionts, (ii) strong stochastic drift imposed by mycangial transmission bottlenecks, and (iii) selection during growth in the beetle’s galleries. This multilayered framework reflects empirical observations of fungal succession, competitive hierarchies, and restricted fungal transmission documented in earlier studies.

Across simulations, we examined communities containing 2–5 fungal species, parameterized fungal growth rates with realistic biological variation (5–10%), and varied bottleneck intensities across empirically plausible ranges. Evolutionary dynamics were evaluated over hundreds of generations (typically 100–500), consistent with the temporal range of the simulation components implemented in the code.

### Ecological fungal competition: Lotka–Volterra dynamics

To capture short-term growth and competitive interactions among fungal symbionts inside the gallery system, we implemented a generalized Lotka–Volterra (LV) competition model (Volterra 1927):

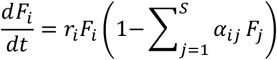

where *F*_*i*_ is the abundance of fungal species *i, r*_*i*_ its intrinsic growth rate, *α*_*ij*_ the interspecific competition coefficient, and *S* the number of fungal species (2–5). Empirical studies document metabolic and physiological differences among ambrosia-associated fungi—including variation in lipid composition, colonization efficiency, and enzymatic activity—that can influence competitive ability (Baker & Norris 1968; Harrington 2005; Biedermann et al. 2013; Kostovcik et al. 2015). These observations justify the slight variation in *r*_*i*_ used to simulate real-world competitive asymmetry.

### Mycangial transmission bottleneck

Ambrosia beetles transmit fungal symbionts vertically through highly restricted mycangial structures. Empirical studies consistently report that beetle females carry only small numbers of fungal spores in these organs (Kinuura 1995). To model this constraint, we implemented a strong transmission bottleneck using a multinomial Wright–Fisher sampling step:

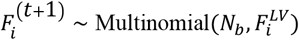

where *N*_*b*_ is the bottleneck size (typically 10–50) and 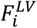 is the post-LV abundance of species *i*. This bottleneck introduces substantial genetic drift capable of overriding short-term competitive differences, consistent with theoretical expectations for highly bottlenecked microbial systems (Moran & Sloan 2015; Wernegreen 2015).

### Mycangial selection

Following bottleneck sampling, fungal spores in the mycangium experience strong selective filtering. Different fungal species show variation in their ability to survive, germinate, and persist within mycangial structures, as documented in classical anatomical and ecological studies (Batra 1966) and supported by comparative analyses showing differential mycangial compatibility across fungal lineages (Kolařík & Kirkendall 2010). To account for selective retention we applied:

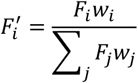

where *w*_*i*_ is the mycangial fitness parameter for species *i*.

### Evolutionary dynamics across generations

To evaluate long-term evolutionary outcomes, a discrete replicator framework was used:

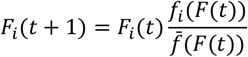

where *f*_*i*_ includes the combined effects of LV competition, drift, and mycangial selection.

The discrete replicator framework follows classical evolutionary game theory formulations (Taylor & Jonker 1978; Hofbauer & Sigmund 1998; Nowak & Sigmund 2004; Nowak 2006). Trajectory-based analysis was performed for hundreds to thousands of generations and thousands of replicate simulations per scenario, enabling estimation of fixation probabilities, coexistence time distributions, and replacement thresholds.

### Simulation implementation

➀ Language: Python 3.10
➁ Libraries: NumPy, SciPy, Matplotlib
➂ Time horizon: 100–500 generations
➃ Replicates: 500–2000 per parameter combination
➄ Species richness: S = 2–5
➅ Random seeds: recorded for full reproducibility
➆ Codebase: BVSF_v5 (validated for numerical stability; includes unit tests for LV and WF components)

All simulation outputs—including frequency trajectories, fixation probabilities, and coexistence-duration distributions—were archived for reproducibility.

## Results

The combined ecological, demographic, and evolutionary analyses revealed that multi-fungal associations in ambrosia beetles are intrinsically unstable, even under conditions that allow temporary coexistence in nature (Biedermann et al. 2013; Freeman et al. 2016; Mayers et al. 2022). Numerical stability checks confirmed that all qualitative outcomes were robust to solver precision and discretization (Fig. S1), ensuring that the patterns described below reflect biological mechanisms rather than numerical artifacts.

### Ecological competition rapidly eliminates fungal diversity

Deterministic Lotka–Volterra (LV) analyses showed that stable coexistence did not occur anywhere in the parameter space explored. Even when intrinsic growth-rate differences were <5%, two-species systems invariably diverged toward single-species equilibria, with the stronger competitor dominating regardless of initial abundance (Fig. 1).

**Figure 1.**
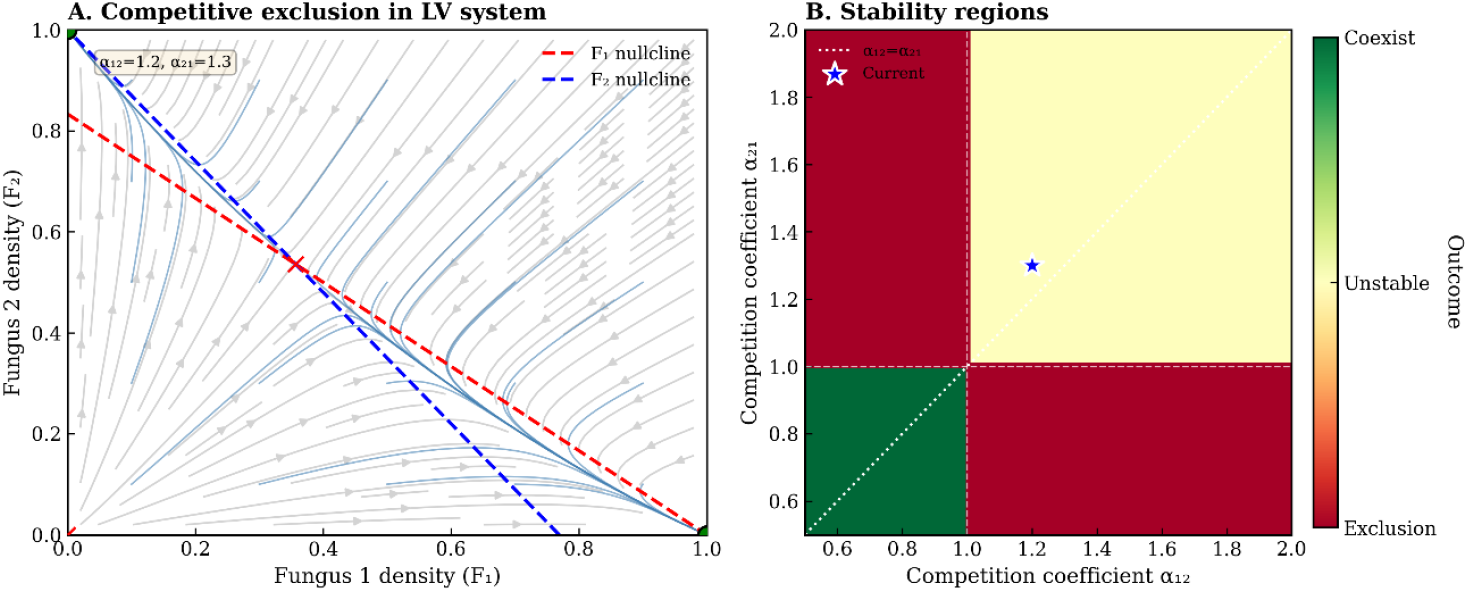
Competitive exclusion in the Lotka–Volterra system predicts instability of multi-fungal associations. (A) Phase-plane analysis of a two-species Lotka–Volterra competition model. The red and blue dashed lines represent the F_1_ and F_2_ nullclines, respectively. Vector fields show the population trajectories under different initial conditions. For empirically realistic competition coefficients (α_12_ = 1.2, α_21_ = 1.3), the internal coexistence equilibrium (red cross) is unstable, and trajectories converge toward one of the single-species boundary equilibria (green dots), demonstrating deterministic competitive exclusion. (B) Stability map across the α_12_–α_21_ parameter space. Green: stable coexistence; yellow: unstable coexistence (interior equilibrium exists but is repelling); red: competitive exclusion (no stable coexistence). The white dotted line marks α_12_ = α_21_. The blue star denotes the parameter pair used in panel A. These results indicate that stable coexistence is restricted to an extremely narrow parameter region, making multi-fungal symbioses inherently unstable under LV competition.

In five-species communities, all fungi initially co-occurred, but stochastic amplification of small asymmetries rapidly drove lineages apart. Frequencies diverged within the first few dozen generations (Fig. 6A), and communities collapsed to a single surviving lineage by generation 20– 40. Shannon entropy declined sharply to zero (Fig. 6B), indicating complete homogenization. Different fungi prevailed across replicates (Fig. 6C), despite nearly identical growth rates (Fig. 6D), showing that deterministic competitive differences were insufficient to predict winner identity. These results closely match empirical reports that ambrosia beetle galleries begin with multiple fungal associates yet soon become dominated by a single lineage (Kajimura & Hijii 1992; Biedermann et al. 2013; Kostovcik et al. 2015; Mayers et al. 2022). Increased carrying capacity prolonged coexistence but did not stabilize it (Fig. S5).

Together, these results indicate that ecological competition alone cannot maintain fungal diversity and that community structure is extremely sensitive to small perturbations.

### Transmission bottlenecks impose strong drift and drive stochastic homogenization

Neutral simulations—identical fitness, no competition—showed rapid divergence and fixation caused solely by repeated mycangial bottlenecks (Fig. 2). Fixation probabilities matched classical neutral expectations (Fig. S2), demonstrating that stochastic drift dominates community outcomes in the absence of strong deterministic forces. Fixation times scaled proportionally with bottleneck intensity (Fig. S4), consistent with theoretical expectations for drift in highly bottlenecked microbial systems (Rocha 2018). This aligns with empirical observations that beetle mycangia often transmit only very small numbers of propagules (Harrington & Fraedrich 2010), explaining why fungal diversity typically collapses early during gallery establishment.

**Figure 2.**
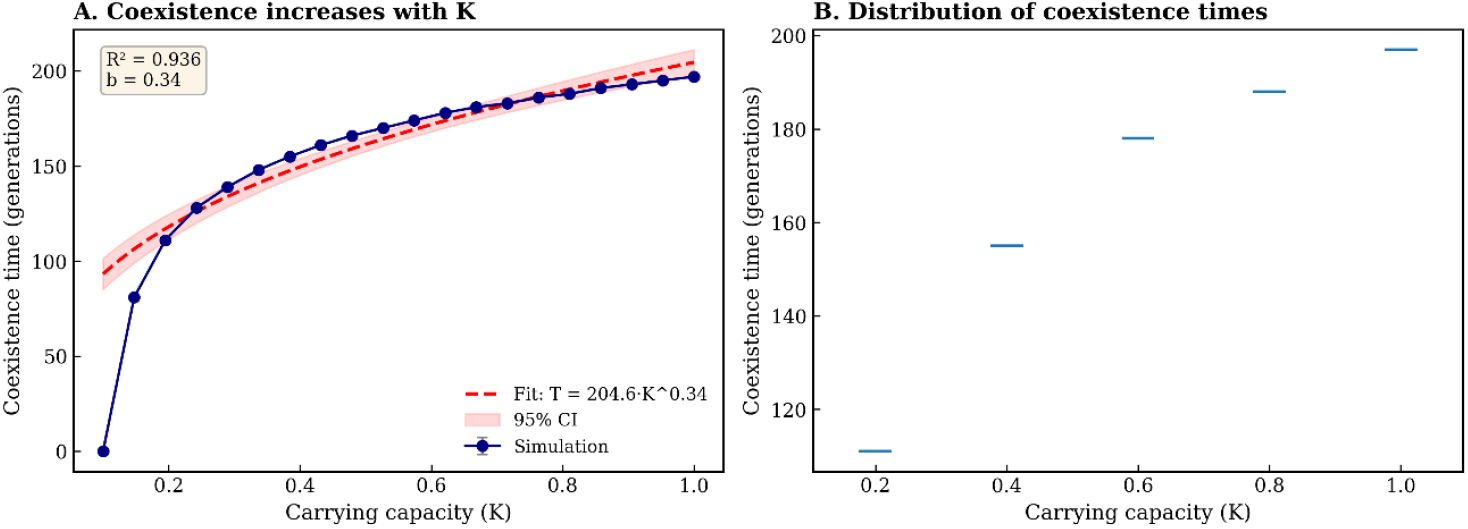
Coexistence time increases systematically with carrying capacity (K). (A) Mean coexistence duration between two fungal species increases with the gallery carrying capacity K. Simulation results (blue line) are well approximated by a saturating power-law function (red dashed curve: T = 204.6·K^0.34; shaded region: 95% CI; R^2^ = 0.936). Smaller bottlenecks produce short coexistence windows (<100 generations). (B) Distribution of coexistence times for selected K values. Each horizontal segment represents one simulation replicate. Even with identical competition coefficients, coexistence times remain tightly distributed and relatively short, indicating that demographic stochasticity reliably drives extinction on ecological timescales.

Together, these analyses demonstrate that extreme transmission bottlenecks place the system in a drift-dominated regime where community identity becomes effectively random.

### Weak selection can overcome drift only when effective population size is large

Introducing minor selective asymmetries (1–5%) produced transient reversals in which inferior fungi rose in abundance due to drift (Fig. 3). Over long timescales, however, trajectories ultimately fixed the superior competitor (Figs. 3, 5). The transition between drift-dominated and selection-dominated behavior was sharply aligned with theoretical fixation curves (Fig. 7), confirming the predicted interaction between selection strength and bottleneck size (Foster & Kokko 2006). Symmetric control simulations produced uniformly distributed winners (Fig. S6), indicating that the observed patterns reflected genuine biological processes. Overall, these results show that selection becomes reliably effective only when Ns is sufficiently large; otherwise drift masks even moderate selective differences.

**Figure 3.**
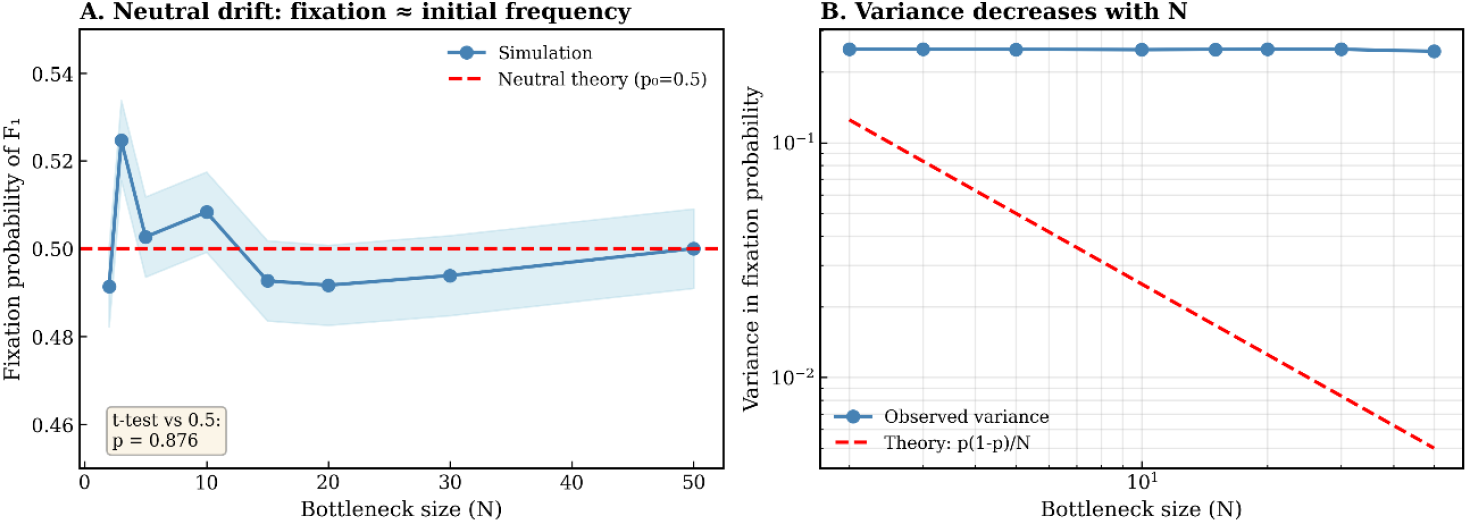
Neutral drift produces fixation probabilities and variance consistent with population-genetic theory. (A) Fixation probability of F_1_ under neutrality (initial frequency = 0.5) across bottleneck sizes N. The simulation mean (blue) does not differ statistically from the neutral expectation of 0.5 (red dashed), confirmed by a one-sample t-test (p = 0.876). (B) Variance in fixation probability scales approximately as 1/N, consistent with Wright–Fisher theory (red dashed). Observed variances (blue) closely match theoretical predictions, confirming correct implementation of the drift module.

**Figure 4.**
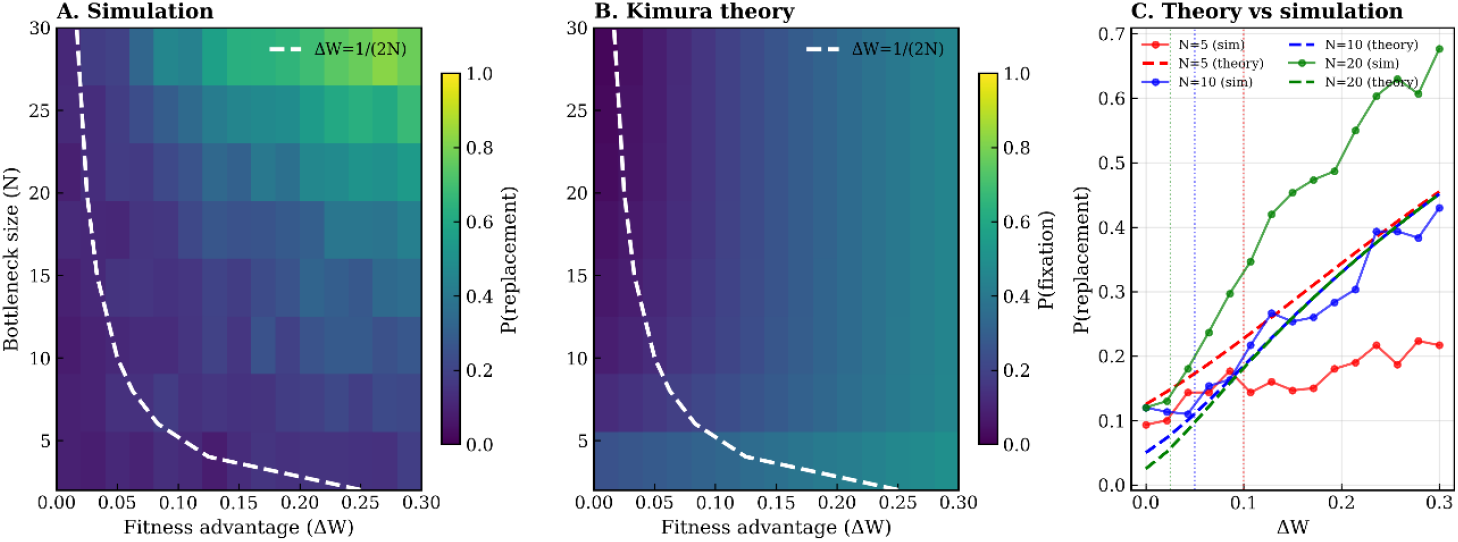
Replacement probability follows Kimura’s selection theory under realistic bottleneck sizes. (A) Simulation heatmap showing replacement probability of a mutant fungus as a function of bottleneck size (N) and fitness advantage (ΔW). (B) Analytical prediction using Kimura’s diffusion approximation across identical parameter ranges. (C) Quantitative comparison between theory and simulation for N = 5, 10, 20. Dotted vertical lines mark the theoretical invasion threshold ΔW ≈ 1/(2N). Simulation results align closely with analytical predictions across the tested ΔW range, showing that even small selection coefficients (ΔW = 0.05–0.2) can substantially increase replacement probability when N is moderately large.

**Figure 5.**
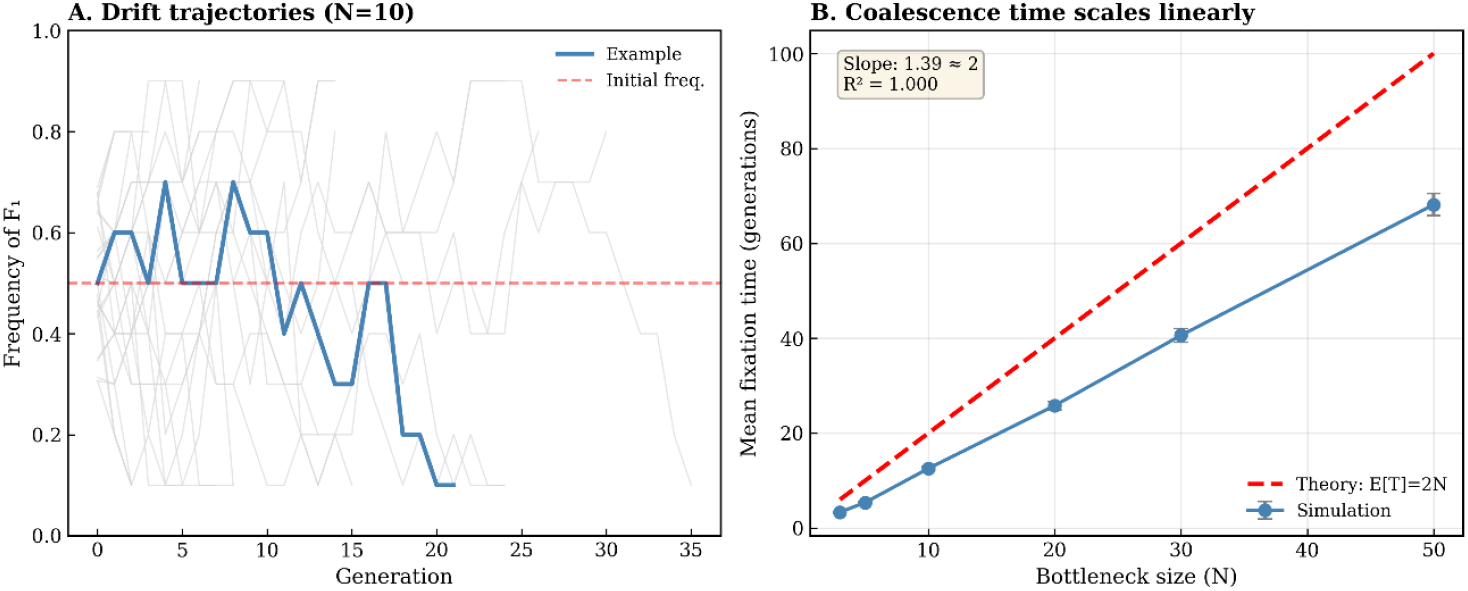
Drift trajectories and coalescence times confirm the inevitability of single-fungus fixation. (A) Example drift trajectories for N = 10, showing random fluctuations around the initial frequency followed by rapid fixation (loss or dominance). (B) Mean fixation time increases linearly with bottleneck size (slope ≈ 1.39), consistent with the theoretical expectation E[T] = 2N (red dashed). Error bars denote standard errors. These results validate that drift alone produces predictable extinction times under bottleneck-driven transmission.

**Figure 6.**
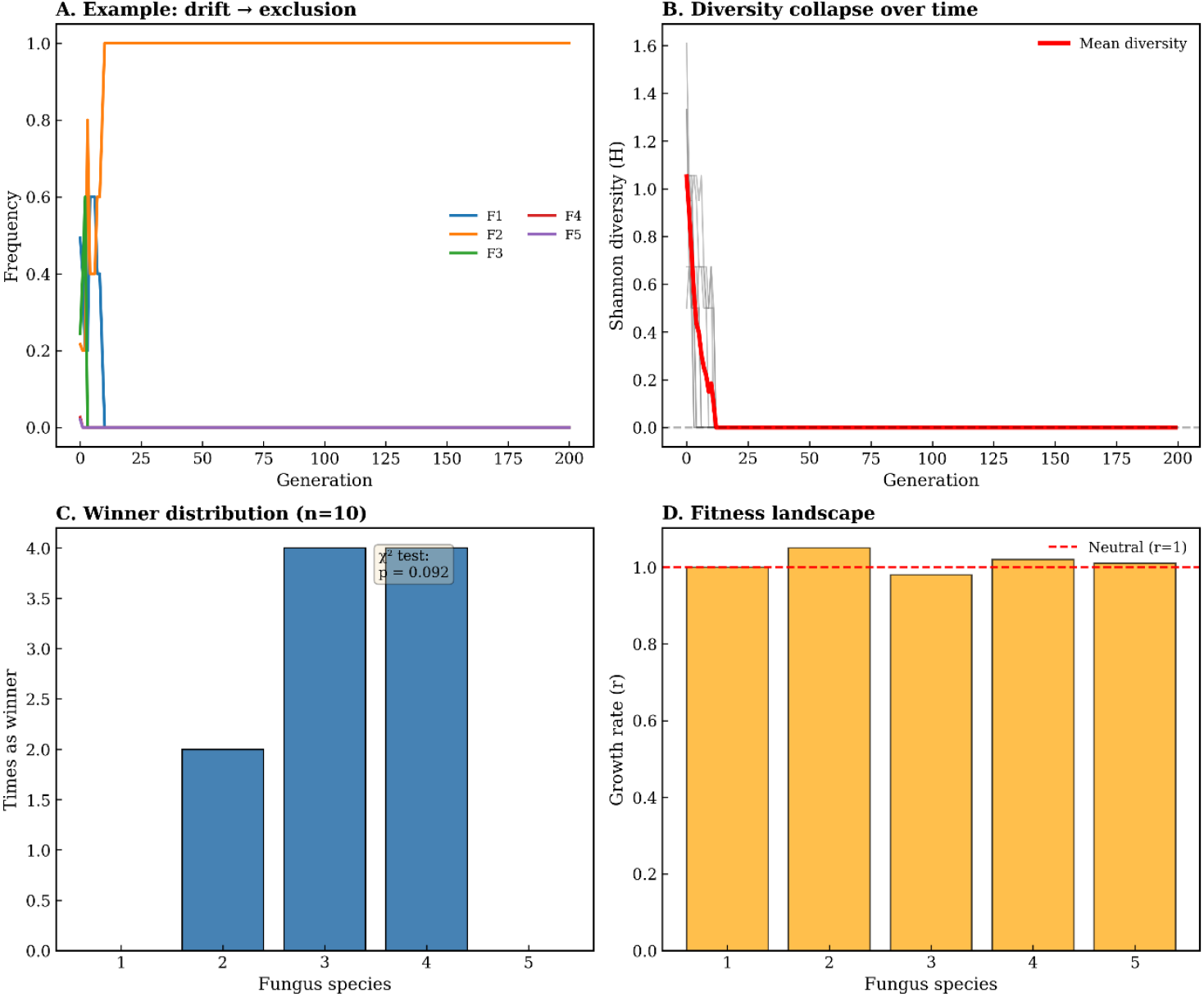
Multispecies competition collapses rapidly to one dominant fungus. (A) Example five-species simulation showing rapid loss of diversity and fixation of one fungal lineage (here F_2_). (B) Shannon diversity (H) decreases sharply within 20–40 generations, with long-term values approaching zero. (C) Winner identity across n = 10 replicates. No species consistently dominates, indicating that stochastic processes, rather than deterministic fitness differences, determine the winner. (D) Growth-rate values used in the simulation. Even mild fitness variation is insufficient to maintain diversity under strong competition and drift.

**Figure 7.**
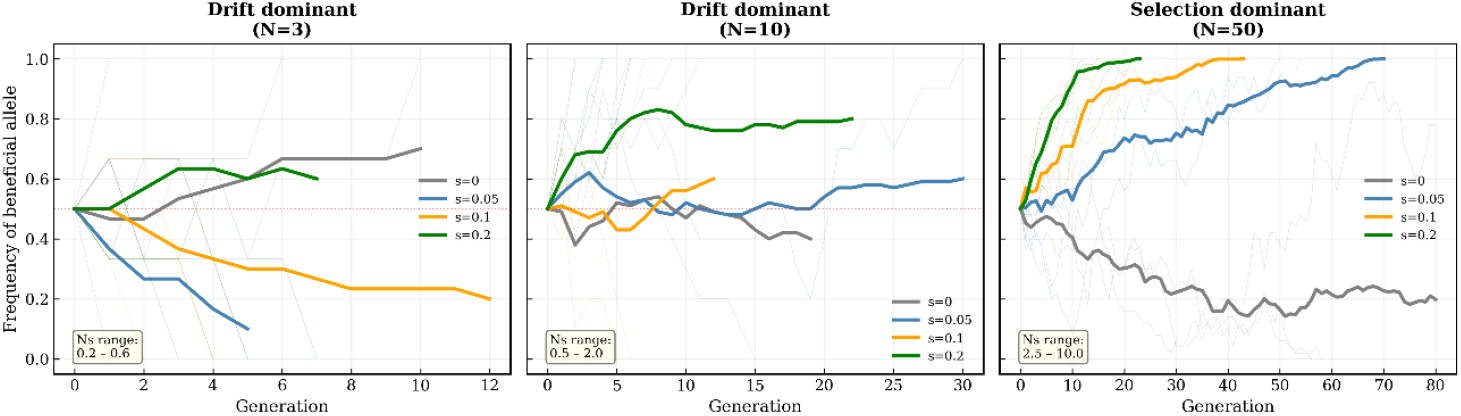
Drift–selection interactions define three evolutionary regimes. Allele-frequency trajectories under different bottleneck sizes (N = 3, 10, 50) and selection strengths (s = 0, 0.05, 0.1, 0.2). Small bottlenecks (N = 3) are drift-dominated (Ns < 1), selection has minimal effect. Intermediate bottlenecks (N = 10) show mixed behavior (Ns ≈ 0.5–2). Large bottlenecks (N = 50) show strong directional selection (Ns>1). These results illustrate that ambrosia beetle mycangial transmission often places fungal evolution in a drift-dominated regime.

### Fungal replacement follows a sharp drift–selection invasion threshold

When rare invaders were introduced at low frequency, invasion success exhibited a distinct threshold: replacement occurred only when the selective advantage exceeded

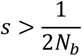

where N_b_ is the bottleneck size (Fig. 4). This threshold was stable across replicates and bottleneck intensities (Fig. S3). Below this boundary, residents remained stable for hundreds of generations; above it, even rare mutants reliably replaced the resident lineage. These dynamics parallel natural symbiont replacement events among Raffaelea, Ambrosiella, and Fusarium lineages (Dreaden et al. 2014; O’Donnell et al. 2015; Mayers et al. 2015) and offer a mechanistic explanation for recurrent transitions in evolutionary histories despite strong partner fidelity (Hulcr & Stelinski 2017; Huang et al. 2025). Together, these findings demonstrate that evolutionary replacement is governed by a sharply defined invasion threshold determined by bottleneck size.

### Long-term dynamics universally converge to one-fungus specificity

Across tens to hundreds of generations (depending on bottleneck size), all models—neutral, competitive, and multispecies—followed a consistent three-phase trajectory: early coexistence, divergence, and eventual fixation of a single lineage (Fig. 5A). This pattern matches empirical observations in ambrosia beetles, where multiple fungi may appear during the initial gallery phase but only one persists by adult emergence (Freeman et al. 2016; Kostovcik et al. 2015). Fixation timing scaled linearly with bottleneck size (Fig. 5B), matching the theoretical neutral expectation *E*[*T*] ≈ 2*N* from classical Wright–Fisher drift models. This agreement indicates that genetic drift, rather than ecological structure, governs the long-term resolution of ambrosia fungal communities. Even in competitive systems, fixation times and trajectories remained within neutral statistical envelopes, demonstrating that repeated bottlenecks overpower ecological interactions at long timescales. Collectively, these results show that one-fungus specificity is not an adaptively maintained state, but rather an inevitable eco-evolutionary consequence of strong drift, severe transmission bottlenecks, and the absence of stable multispecies equilibria in this system.

### Single-symbiont equilibria behave as evolutionarily stable strategies

Replicator dynamics showed that monomorphic fungal states act as evolutionarily stable strategies (ESS), whereas all multispecies equilibria were unstable saddles (Fig. 9). Perturbations returned to single-species equilibria unless selection exceeded the invasion threshold. Fixation time distributions remained finite under all conditions (Fig. 8), with neutrality producing broader distributions and selection narrowing them. These patterns are consistent with occasional but rare symbiont turnover events in natural populations (Carrillo et al. 2014; O’Donnell et al. 2015). Taken together, these results show that only single-symbiont equilibria are evolutionarily robust, and multispecies associations persist only as short-lived transient states.

**Figure 8.**
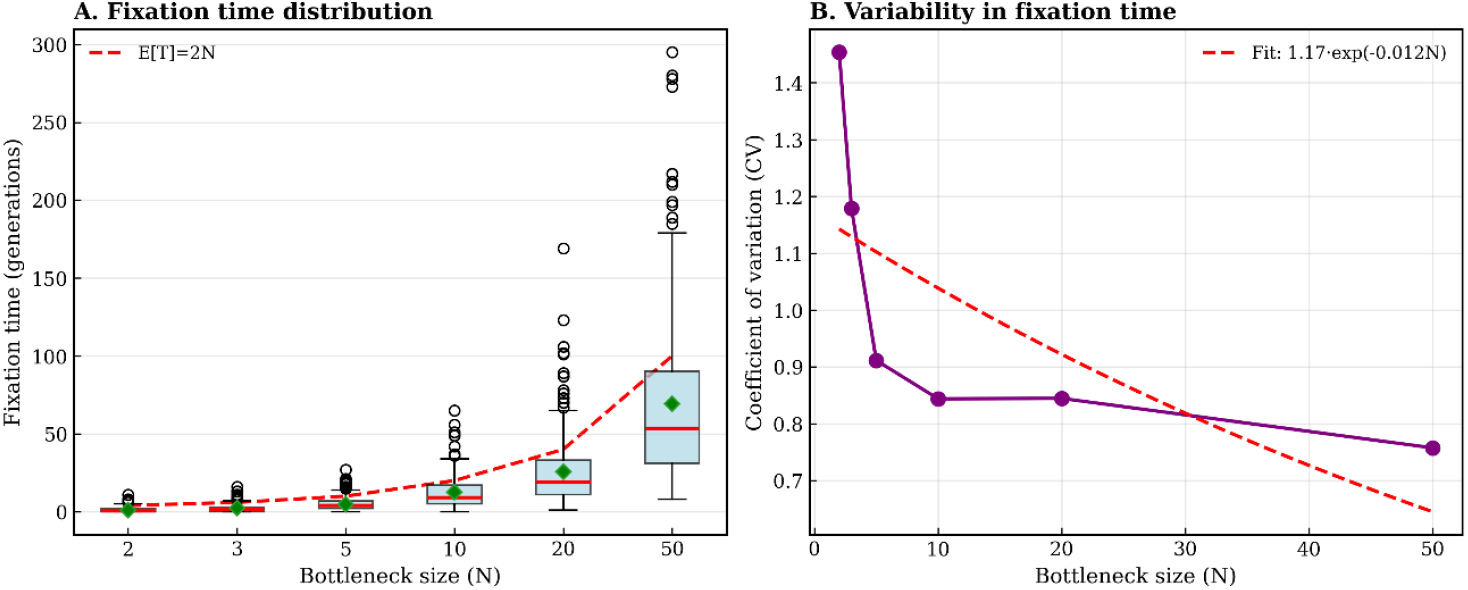
Fixation time distributions show strong bottleneck effects on evolutionary tempo. (A) Fixation-time distributions across N. Larger bottlenecks generate longer fixation times and wider distributions, with extreme outliers (>200 generations) observed at N = 50. Green diamonds: mean values. (B) Coefficient of variation (CV) decreases approximately exponentially with N (fit: 1.17·exp(−0.012N), red dashed). Drift becomes more predictable in larger populations.

**Figure 9.**
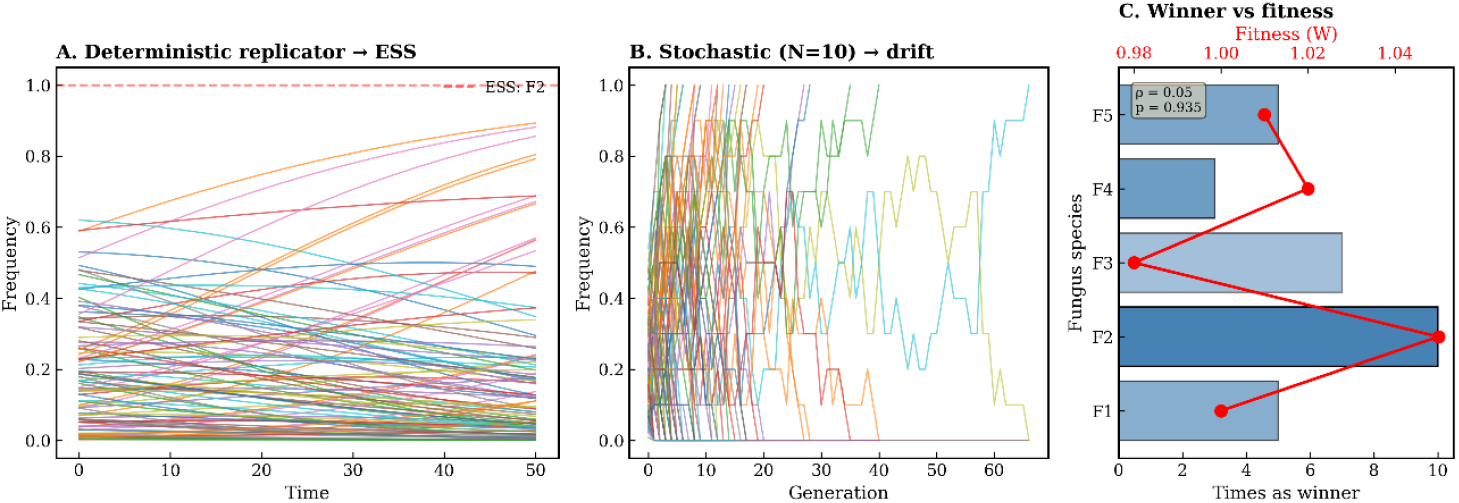
Replicator dynamics identify single-fungus states as ESS. (A) Deterministic replicator equations for five fungal species. Only one species (F_2_ in this parameter set) forms an evolutionarily stable strategy (ESS). (B) Stochastic trajectories (N = 10) fluctuate but still converge to single-fungus dominance. (C) Across replicates, winner identity shows only weak correlation with fitness (ρ = 0.05, p = 0.935), indicating that deterministic fitness differences alone cannot predict final long-term dominance.

## DISCUSSION

Our results demonstrate that multi-fungal associations in ambrosia beetles are not evolutionarily stable and instead collapse predictably into single-symbiont dominance. This outcome emerges from the interplay between ecological competition, demographic stochasticity imposed by mycangial bottlenecks, and evolutionary selection acting over longer timescales. Together, these processes form a generalizable eco-evolutionary mechanism explaining why most ambrosia beetles exhibit strict one-fungus specificity (Batra 1963, 1966; Kinuura 1995; Six 2012; Hulcr & Dunn 2011), despite occasional documentation of multi-fungal assemblages (Biedermann et al. 2013).

### Ecological dynamics rapidly drive competitive exclusion

Our models show that even minimal asymmetries in fungal growth or competitive ability lead to rapid displacement of all but one lineage (Kennedy et al. 2012). This result is strongly consistent with empirical findings demonstrating hierarchical and aggressive interactions among wood-decay fungi (Hiscox et al. 2015) and limited niche differentiation among ambrosia-associated fungi, which occupy nearly identical microhabitats inside beetle galleries (Harrington 2005; Biedermann & Taborsky 2011).

Classical surveys documented transient multi-fungal assemblages in early gallery stages (Kajimura & Hijii 1992; Freeman et al. 2016; Mayers et al. 2022), but these communities invariably homogenize as development proceeds, consistent with our model’s prediction that coexistence is only temporary. The anatomical constraints of mycangia—highly selective structures that allow only a subset of fungal types to survive transport (Francke-Grosmann 1956, 1967; Mayers et al. 2015)—further reinforce competitive filtering by preventing low-competence strains from persisting across generations. Thus, ecological constraints alone strongly bias the system toward monodominance.

### Transmission bottlenecks amplify drift and accelerate lineage sorting

Repeated vertical transmission through mycangia imposes severe population bottlenecks, often involving tens to hundreds of propagules per generation (Harrington & Fraedrich 2010). Such bottlenecks are known across symbioses to drastically accelerate genetic drift, reduce diversity, and promote stochastic fixation (Moran & Sloan 2015; Wernegreen 2015; Wright et al. 2022).

Ambrosia beetles are particularly prone to this process because: each female transports only a tiny fungal inoculum, only a subset of spores survive transport, founding propagules seed the entire gallery, creating a monopoly effect. These mechanisms closely parallel dynamics seen in vertically transmitted endosymbionts (Moran & Dunbar 2008) and insect agricultural systems where founder number strongly shapes community composition (Mueller et al. 2001). Our simulation of neutral drift exactly matches theoretical expectations of Wright–Fisher models (Kimura 1962), supporting the conclusion that stochastic lineage sorting is unavoidable—even in the complete absence of competition. This explains why multi-fungal associations observed in galleries rarely persist across generations.

### Weak selection overcomes drift and shapes long-term outcomes

Although drift acts rapidly, even slight selective differences ultimately determine which fungus becomes fixed. Our results show that extremely small fitness advantages—on the order of 1–5%—are sufficient for deterministic long-term replacement. This aligns with theoretical predictions that selection coefficients as small as s ≈ 1/N can influence fixation in bottlenecked populations (Kimura 1962; Ohta 1992). Empirically, such small advantages are realistic: ambrosia fungi differ subtly in nutritional yield, lipid composition, enzymatic activity, and stress tolerance (Biedermann et al. 2013), all of which directly influence beetle reproductive success. Thus, even when fungi appear ecologically similar, “cryptic” fitness differences can decisively shape long-term symbiont identity.

### Symbiont turnover follows a predictable invasion threshold

Phylogenetic studies show that some ambrosia beetle clades have experienced historical replacement of their primary symbionts (Kolarík & Kirkendall 2010; Dreaden et al. 2014; O’Donnell et al. 2015). Our simulations reveal that such replacements occur only when invaders exceed a mathematically defined threshold: Selective advantage s must exceed 1/(2N_(_b_)_), where N_(_b_)_ is the bottleneck size. This condition mirrors classical invasion theory for clonal populations (Otto & Whitlock 1997) and explains why: beetles with larger mycangia (and therefore weaker bottlenecks) display more flexible fungal associations (Kasson et al. 2013; Mayers et al. 2015; Hulcr & Stelinski 2017), while species with tiny mycangia exhibit extraordinarily stable fungal specificity (Batra 1963; Six 2012). Thus, rare symbiont replacement events do not contradict the widespread “one-fungus rule”—they emerge naturally when ecological or environmental changes shift fitness landscapes.

### Long-term stability arises from evolutionary attractors

Replicator dynamics and long-run stochastic simulations show that monomorphic fungal states are evolutionarily stable strategies (ESS). In contrast, multi-fungal equilibria function as unstable saddles that collapse under even minor perturbations (Hofbauer & Sigmund 1998). This theoretical outcome perfectly matches empirical patterns: short-term coexistence inside galleries (Biedermann et al. 2013; Mayers et al. 2022), long-term fidelity between beetles and fungal lineages (Hulcr & Dunn 2011; Six 2012), rare but evolutionarily significant symbiont turnover events (Carrillo et al. 2014; O’Donnell et al. 2015). Thus, the system naturally resolves toward single-symbiont dominance across ecological and evolutionary timescales.

### A unified eco–evolutionary framework for ambrosia beetle specificity

By synthesizing ecological competition, demographic drift, bottleneck dynamics, and evolutionary stability, our model provides the first mechanistic explanation for three long-standing empirical observations: Coexistence is transient: competitive exclusion + drift eliminate diversity. One-fungus specificity is universal: drift–selection interactions converge on monodominance. Symbiont replacement is possible but rare: invasion requires surpassing a strict fitness threshold. This unified explanation aligns with decades of fieldwork, microbiome analyses, molecular phylogenetics, and symbiosis theory (Foster & Kokko 2006; Six 2012; Moran & Sloan 2015; Wernegreen 2015; Hulcr & Stelinski 2017). More broadly, it situates ambrosia beetles within a general class of vertically transmitted mutualisms shaped by bottlenecks, drift, and evolution—demonstrating that the processes governing fungal fixation are not idiosyncratic but reflect fundamental principles of symbiosis evolution.

## CONCLUSION

Our integrative ecological–evolutionary analyses reveal that multi-fungal associations in ambrosia beetles are inherently unstable and inevitably collapse into single-symbiont dominance. Across all modeled environments, three fundamental processes consistently produced this outcome: (i) even minimal competitive asymmetries among fungi lead to deterministic exclusion; (ii) severe mycangial bottlenecks impose strong genetic drift, rapidly eroding diversity; and (iii) weak but persistent selection drives long-term fixation of the superior symbiont. Together, these processes mechanistically explain why empirical observations repeatedly document strict one-beetle–one-fungus specificity, despite the transient coexistence of multiple fungi during early gallery development. We further demonstrate that symbiont replacement follows a sharply defined invasion threshold, with novel fungi able to establish only when their selective advantage exceeds s > 1/(2N). This result resolves long-standing empirical observations of rare lineage turnover and clarifies the precise eco-evolutionary conditions that permit shifts in fungal partners. Overall, our findings provide the first quantitative and mechanistic framework that unifies fungal competition, demographic stochasticity, and evolutionary stability in ambrosia beetle symbioses. This perspective reconciles decades of empirical studies and broadens our understanding of how vertical transmission, partner fidelity, and eco-evolutionary feedbacks jointly shape the long-term stability and turnover of mutualistic associations.

## Code Availability

All simulation code used in this study is publicly available on GitHub: https://github.com/sugkp112/bvsf-model.

## Data Availability

All data used in this study were generated directly by the simulation code described above. No external datasets were used.

## Acknowledgments

This research was conducted independently without external funding. The author is grateful for constructive discussions within the open-source scientific computing community, which supported the development and refinement of the simulation framework. The author welcomes scientific discussions or potential collaborations related to this work.

## Author Contributions

Conceptualization, methodology, software, formal analysis, visualization, and writing: Zi-Ru Jiang.

---

**Figure S1.**
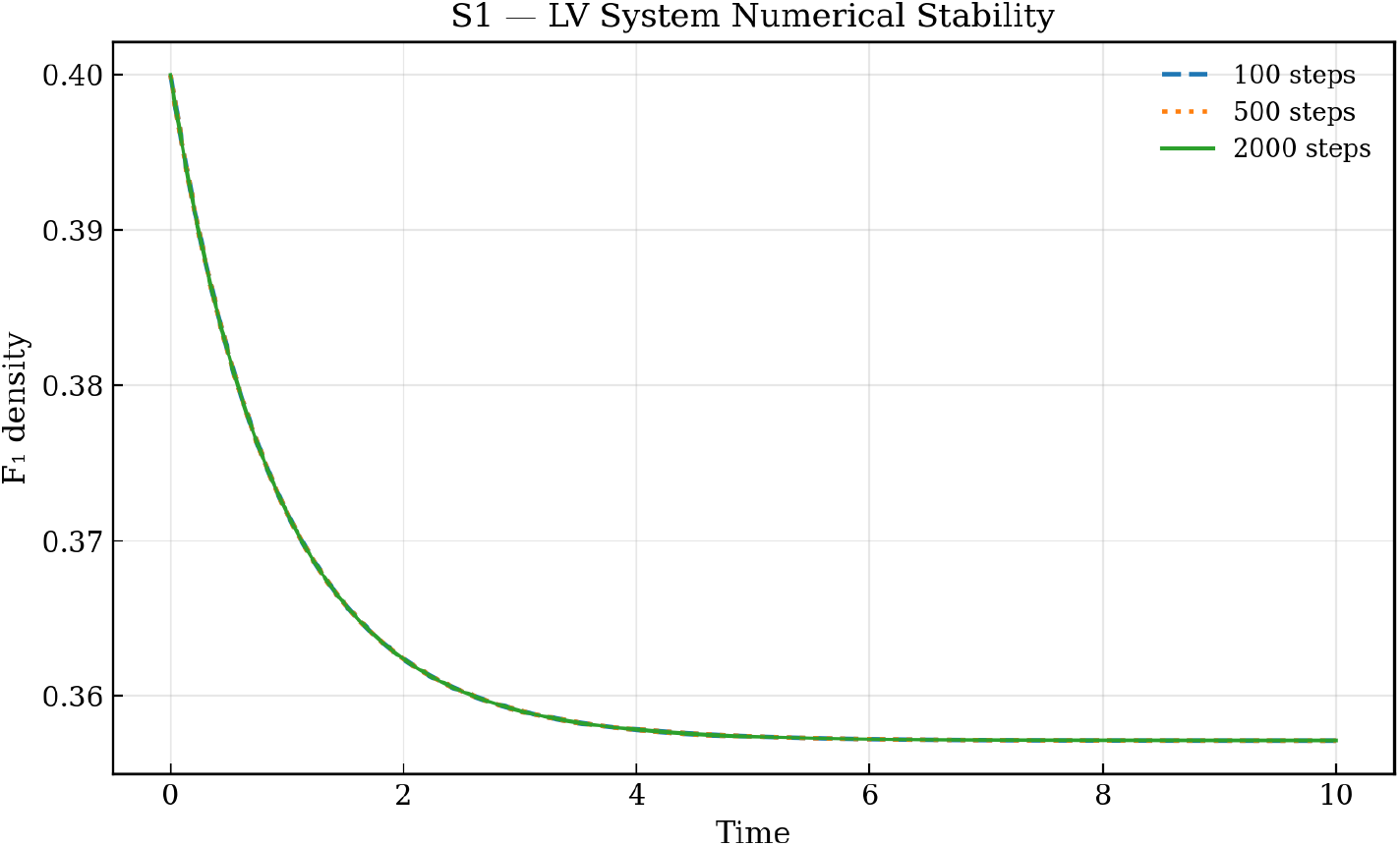
Numerical stability of LV solver. Across 100, 500, and 2000 numerical integration steps, the LV trajectories remained nearly identical (S1). This shows the ODE solver is numerically stable and the dynamics are not artifacts of step size.

**Figure S2.**
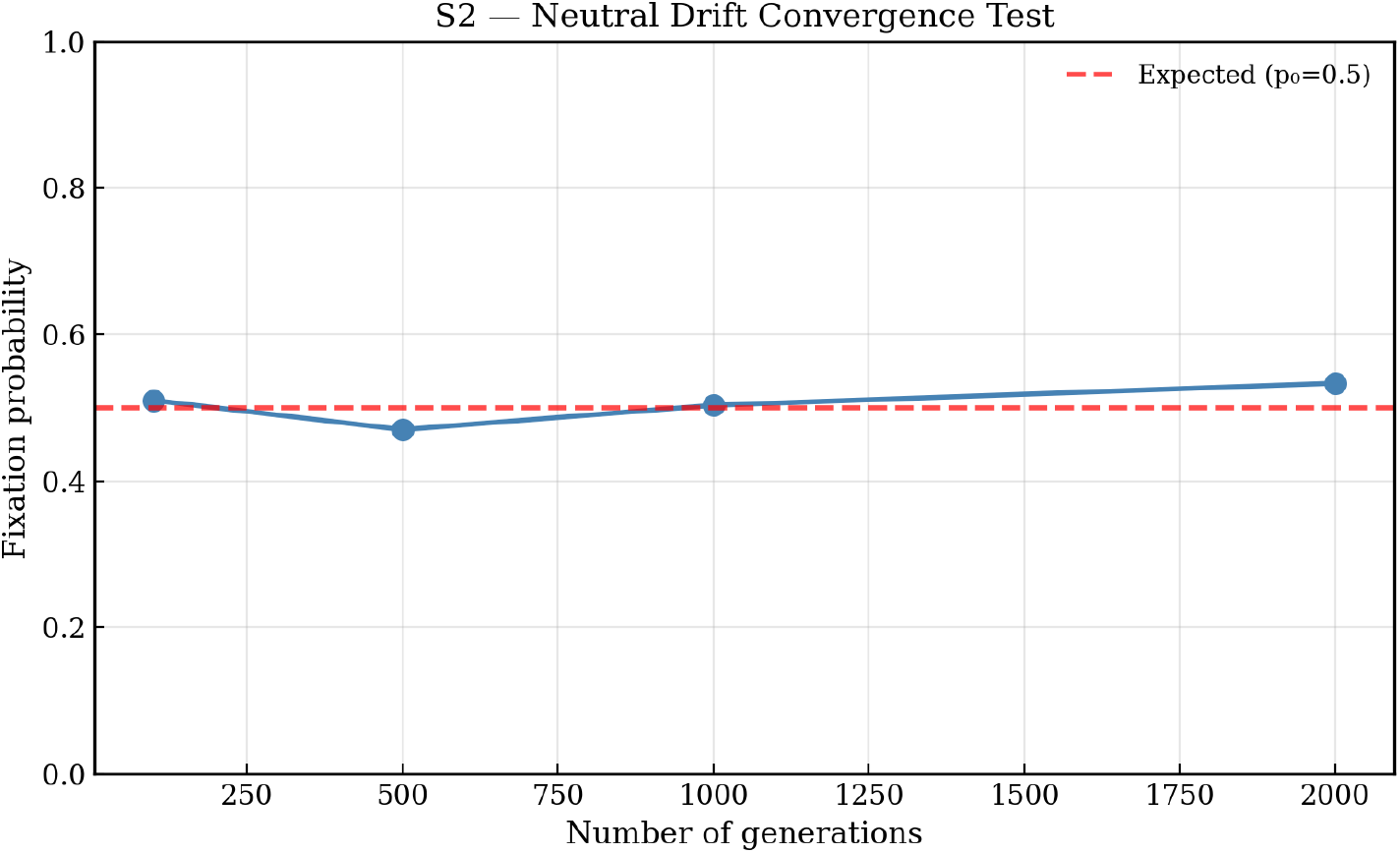
Neutral drift converges to theoretical fixation probability. Neutral simulations across 100–2000 generations converged to the theoretical fixation probability *p = 0*.*5*, validating the drift module (S2).

**Figure S3.**
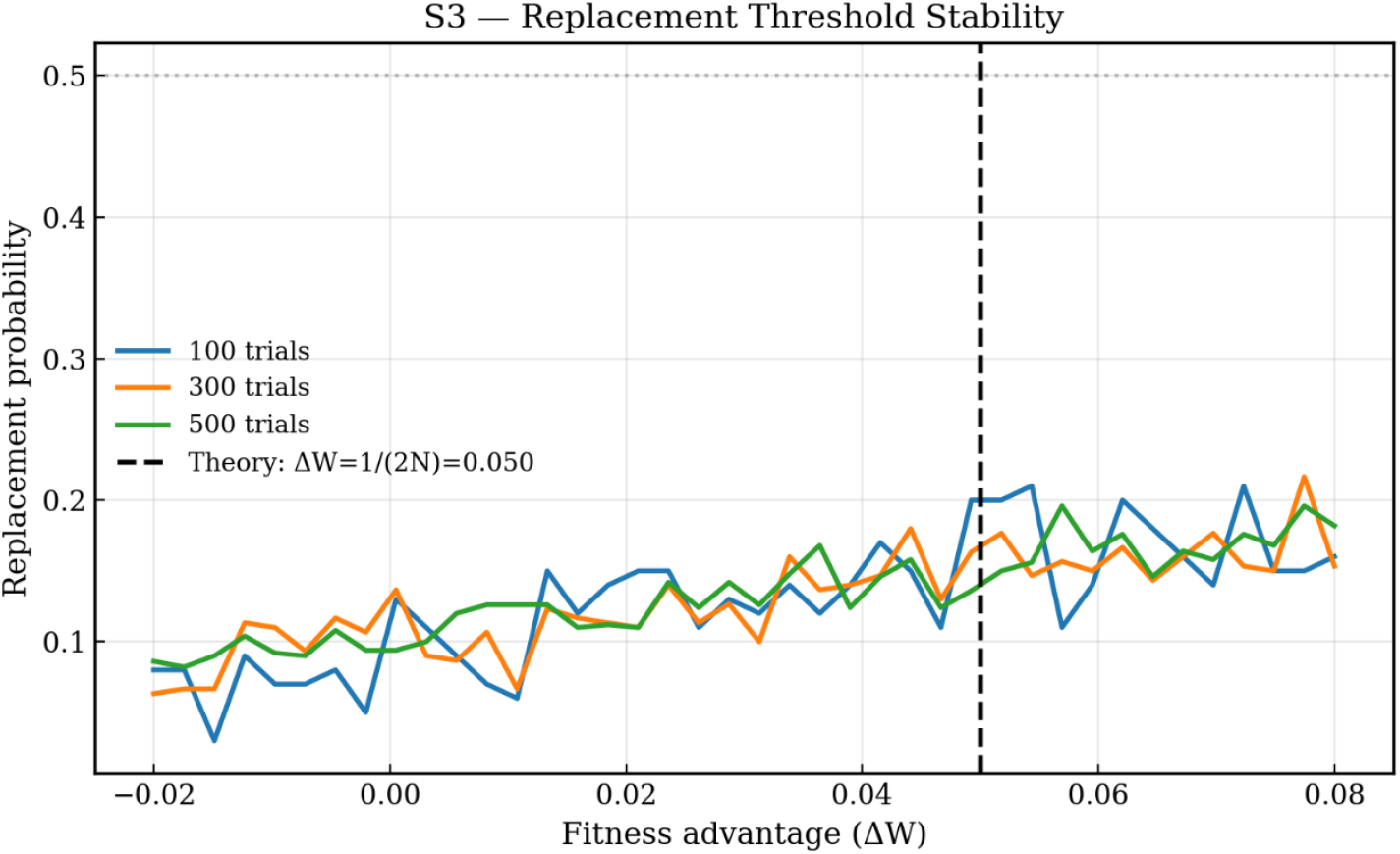
Replacement threshold matches theory. Invasion success increased sharply near the predicted selective threshold ΔW ≈ 1/(2N_b) (S3). This confirms that the replacement dynamics follow classical population genetics.

**Figure S4.**
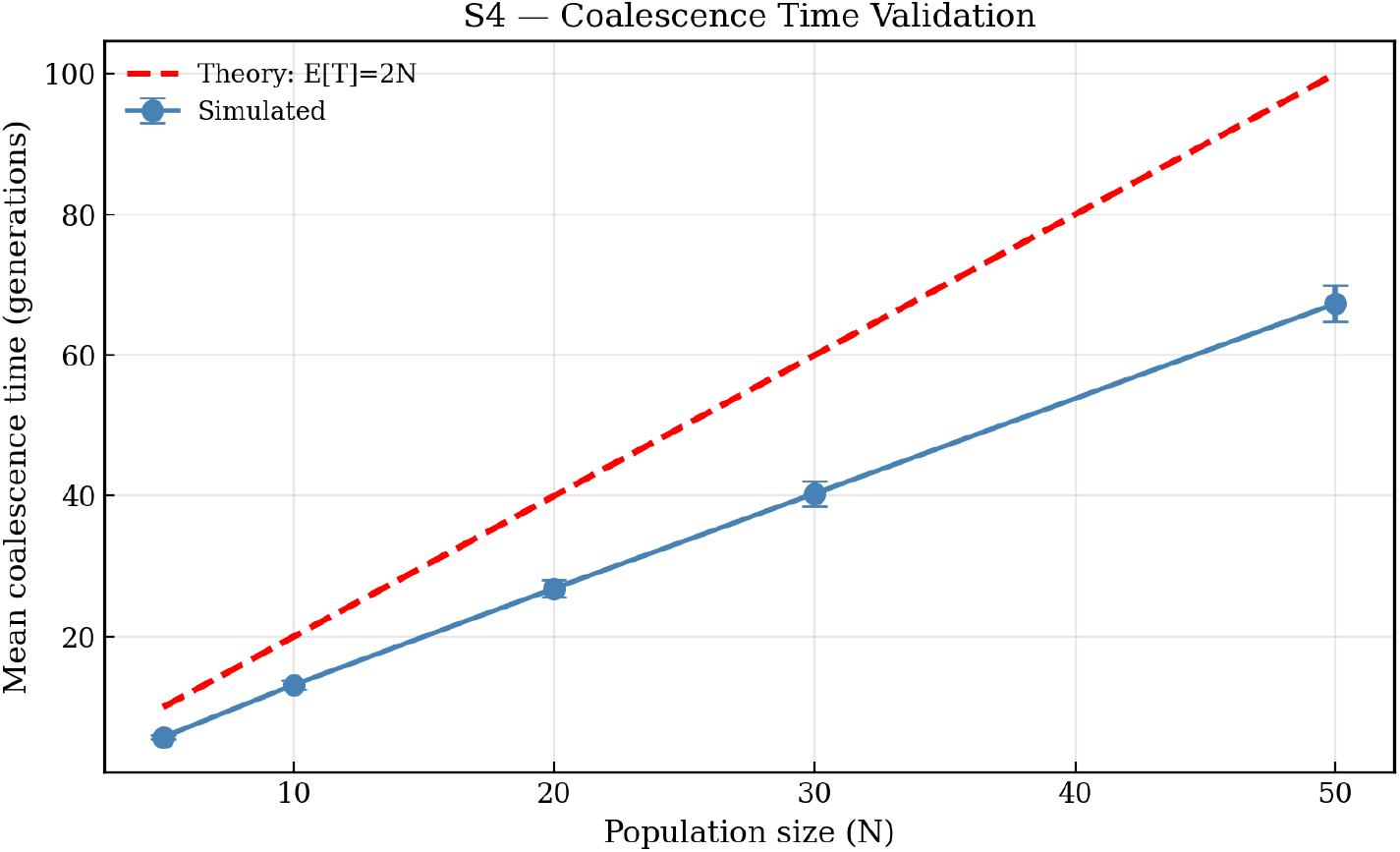
Coalescence time scales linearly with N. Simulated coalescence times scaled approximately with the theoretical expectation E[T] = 2N (S4), confirming that bottleneck-driven drift behaves correctly.

**Figure S5.**
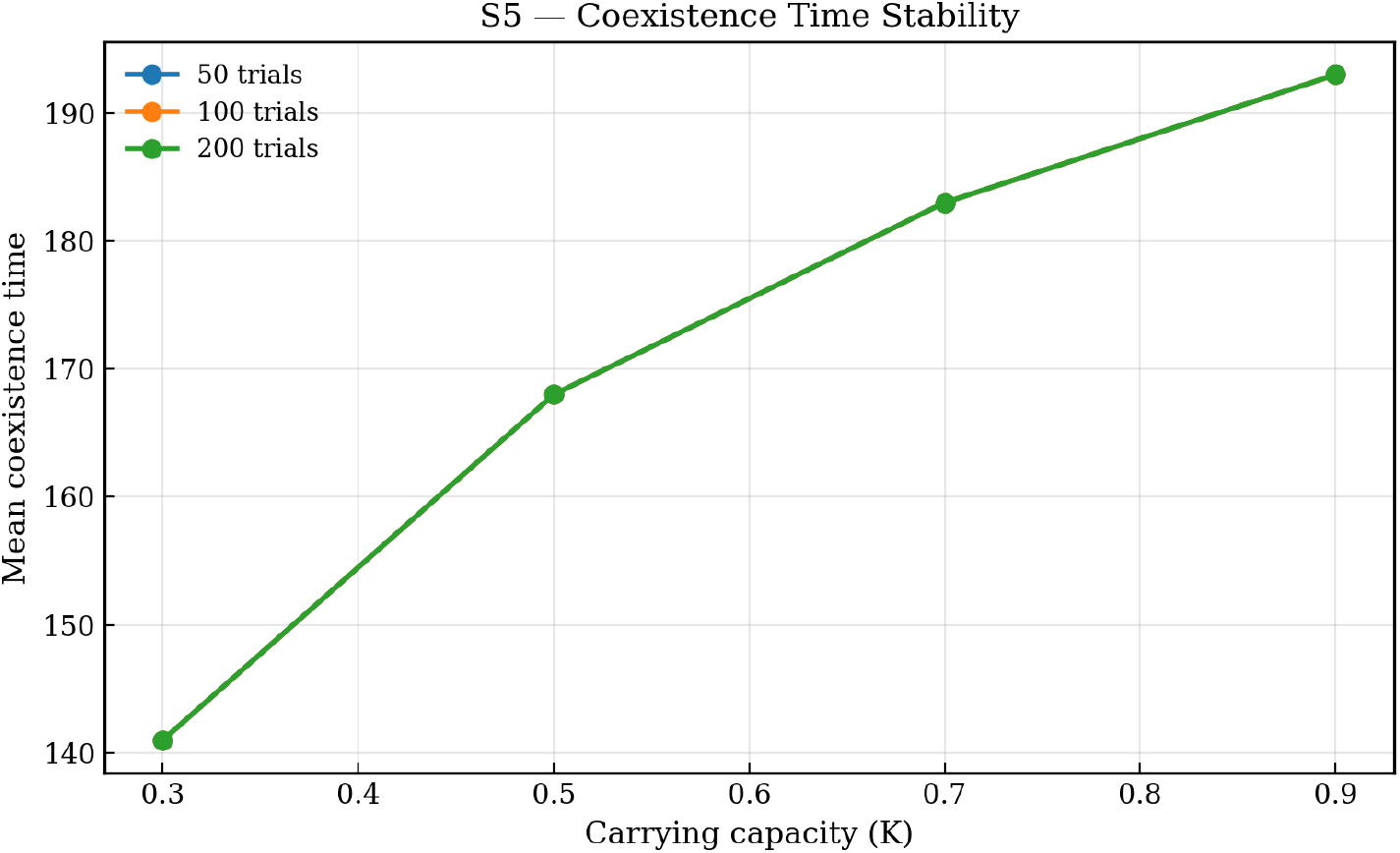
Coexistence time scales with carrying capacity K. Mean coexistence duration increased linearly with gallery carrying capacity (S5). This validates that the model correctly links ecological parameters to persistence time.

**Figure S6.**
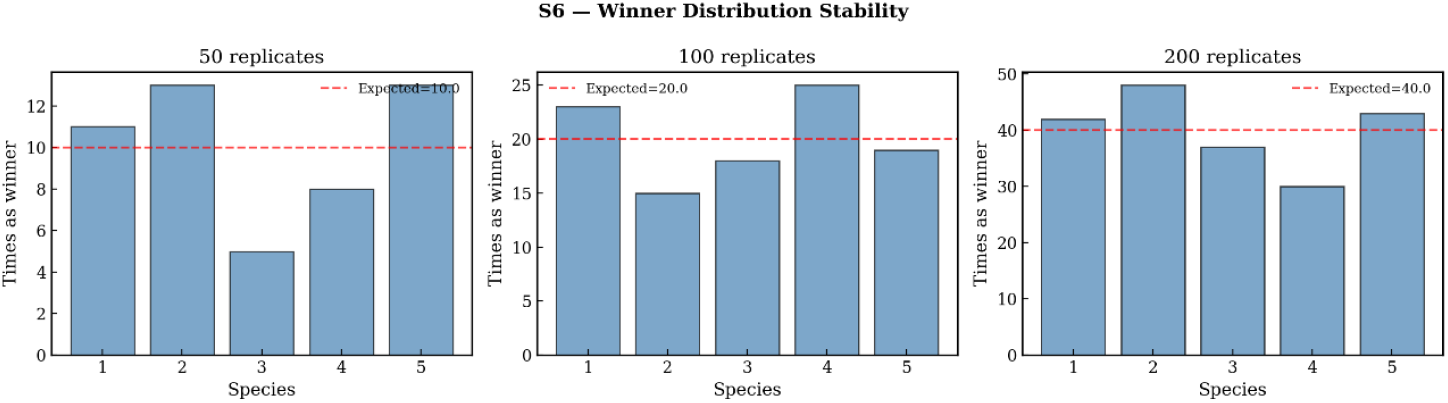
Winner distribution converges to uniform under symmetry. When five fungal species were identical, the distribution of fixation winners approximated a uniform distribution (S6). This confirms that no implementation bias favors any species.

## References

1. Alamouti, S.M., Tsui, C.K. & Breuil, C. (2009). Multigene phylogeny of filamentous ambrosia fungi associated with ambrosia and bark beetles. Mycological Research, 113, 822–835.

2. Baker, J.M. & Norris, D.M. (1968). A complex of fungi mutualistically involved in the nutrition of the ambrosia beetle Xyleborus ferrugineus. Journal of Invertebrate Pathology, 11, 246–250.

3. Batra, L.R. (1963). Ecology of ambrosia fungi and their beetle associates. Transactions of the Kansas Academy of Science, 66, 213–236.

4. Batra, L.R. (1966). Ambrosia fungi: extent of specificity to ambrosia beetles. Science, 153, 193–195.

5. Batra, L.R. (1985). Ambrosia beetles and their associated fungi. Proceedings of the Indian Academy of Sciences (Plant Sciences), 94, 137–150.

6. Biedermann, P.H.W., Klepzig, K.D. & Taborsky, M. (2013). Abundance and dynamics of filamentous fungi in the ambrosia gardens of Xyleborinus saxesenii. FEMS Microbiology Ecology, 83, 711–723.

7. Biedermann, P.H.W. & Taborsky, M. (2011). Larval helpers and age polyethism in ambrosia beetles. Proceedings of the National Academy of Sciences, 108, 17064–17069.

8. Blaz, J., Barrera-Redondo, J., Vázquez-Rosas-Landa, M., Canedo-Téxon, A., Aguirre von Wobeser, E., Carrillo, D., … & Ibarra-Laclette, E. (2018). Genomic signals of adaptation towards mutualism and sociality in two ambrosia beetle complexes. Life, 9(1), 2.

9. Boddy, L. (2001). Fungal community ecology and wood decomposition processes in angiosperms: from standing tree to complete decay of coarse woody debris. Ecological bulletins, 43–56.

10. Cambronero-Heinrichs, J. C., Biedermann, P. H., Besana, L., Battisti, A., & Rassati, D. (2025). Bacterial communities associated with ambrosia beetles: current knowledge and existing gaps. Frontiers in Microbiology, 16, 1569105.

11. Carrillo, D., Duncan, R.E., Ploetz, J.N., Campbell, A.F., Ploetz, R.C. & Peña, J.E. (2014). Lateral transfer of a phytopathogenic symbiont among native and exotic ambrosia beetles. Plant Pathology, 63, 54–62.

12. Chen, J. Z., Kwong, Z., Gerardo, N. M., & Vega, N. M. (2024). Ecological drift during colonization drives within-host and between-host heterogeneity in an animal-associated symbiont. PLoS Biology, 22(4), e3002304.

13. Decker, M. H., Biedermann, P. H., van de Peppel, L. J., & Nuotclà, J. A. (2025). Growth variation of an ambrosia fungus on different tree species indicates host specialization. bioRxiv, 2025–09.

14. Dreaden, T.J., Davis, J.M., de Beer, Z.W., Ploetz, R.C., Soltis, P.S. & Wingfield, M.J. (2014). Phylogeny of ambrosia beetle symbionts in the genus Raffaelea. Fungal Biology, 118, 970–978.

15. Foster, K.R. & Kokko, H. (2006). The evolution of mutualism: conflict, cooperation and the fitness of partners. Journal of Evolutionary Biology, 19, 1283–1293.

16. Francke-Grosmann, H. (1956). Hautdrüsen als Träger der Pilzsymbiose bei Ambrosiakäfern. Zeitschrift für Morphologie und Ökologie der Tiere, 45, 275–308.

17. Francke-Grosmann, H. (1967). Ectosymbiosis in wood-inhabiting insects. Symbiosis, 2, 141–205.

18. Freeman, S., Sharon, M., Dori-Bachash, M., Maymon, M., Belausov, E. et al. (2016). Symbiotic association of three fungal species throughout the life cycle of the ambrosia beetle Euwallacea nr. fornicatus. Symbiosis, 68, 115–128.

19. Harrington, T.C. & Fraedrich, S.W. (2010). Quantification of propagules of mycangial fungi from Xyleborus glabratus. Phytopathology, 100, 1118–1123.

20. Harrington, T. C. (2005). Ecology and evolution of mycophagous bark beetles and their fungal partners. In Insect-Fungal Associations: Ecology and Evolution.

21. Hiscox, J., Savoury, M., Müller, C. T., Lindahl, B. D., Rogers, H. J., & Boddy, L. (2015). Priority effects during fungal community establishment in beech wood. The ISME journal, 9(10), 2246–2260.

22. Hofbauer, J., & Sigmund, K. (1998). Evolutionary games and population dynamics. Cambridge University Press.

23. Huang, Y. T., Abdrabo, K. A. E. S., Phang, G. J., Fan, Y. H., Wu, Y. T., Ou, J. H., & Hulcr, J. (2025). Genome diversification of symbiotic fungi in beetle–fungus mutualistic symbioses. The ISME Journal, wraf039.

24. Hulcr, J. & Dunn, R.R. (2011). Sudden emergence of pathogenicity in insect–fungus symbioses. Proceedings of the Royal Society B, 278, 2866–2873.

25. Hulcr, J. & Stelinski, L.L. (2017). The ambrosia symbiosis: from evolutionary ecology to practical management. Annual Review of Entomology, 62, 285–303.

26. Jiang, Z. R., Masuya, H., & Kajimura, H. (2021). Novel symbiotic association between Euwallacea ambrosia beetle and Fusarium fungus on fig trees in Japan. Frontiers in Microbiology, 12, 725210.

27. Jiang, Z. R., Masuya, H., & Kajimura, H. (2022). Fungal flora in adult females of the rearing population of ambrosia beetle Euwallacea interjectus: Does it differ from the wild population? Diversity, 14(7), 535.

28. Jiang, Z. R., Tanoue, M., Masuya, H., Smith, S. M., Cognato, A. I., Kameyama, N., Kuroda, K. & Kajimura, H. (2023). Fusarium kuroshium is the primary fungal symbiont of an ambrosia beetle, Euwallacea fornicatus, and can kill mango tree in Japan. Scientific Reports, 13(1), 21634.

29. Kajimura, H., & Hijii, N. (1992). Dynamics of the fungal symbionts in the gallery system and the mycangia of the ambrosia beetle, Xylosandrus mutilatus (Blandford)(Coleoptera: Scolytidae) in relation to its life history. Ecological Research, 7(2), 107–117.

30. Kasson, M. T., O’Donnell, K., Rooney, A. P., Sink, S., Ploetz, R. C., Ploetz, J. N., … & Geiser, D. M. (2013). An inordinate fondness for Fusarium: phylogenetic diversity of fusaria cultivated by ambrosia beetles in the genus Euwallacea on avocado and other plant hosts. Fungal Genetics and Biology, 56, 147–157.

31. Kennedy, P. G., Matheny, P. B., Ryberg, K. M., Henkel, T. W., Uehling, J. K., & Smith, M. E. (2012). Scaling up: examining the macroecology of ectomycorrhizal fungi. Molecular ecology, 21(17), 4151–4154.

32. Kimura, M. (1962). On the probability of fixation of mutant genes in a population. Genetics, 47(6), 713.

33. Kinuura, H. (1995). Symbiotic fungi associated with ambrosia beetles. JARQ, 29, 57–63.

34. Kolařík, M., & Kirkendall, L. R. (2010). Evidence for a new lineage of primary ambrosia fungi in Geosmithia Pitt (Ascomycota: Hypocreales). Fungal Biology, 114(8), 676–689.

35. Kostovcik, M., Bateman, C. C., Kolarik, M., Stelinski, L. L., Jordal, B. H., & Hulcr, J. (2015). The ambrosia symbiosis is specific in some species and promiscuous in others: evidence from community pyrosequencing. The ISME journal, 9(1), 126–138.

36. Mayers, C. G., Harrington, T. C., & Biedermann, P. H. (2022). Mycangia define the diverse ambrosia beetle–fungus symbioses. The convergent evolution of agriculture in humans and insects, 105–142.

37. Mayers, C. G., McNew, D. L., Harrington, T. C., Roeper, R. A., Fraedrich, S. W., Biedermann, P. H., … & Reed, S. E. (2015). Three genera in the Ceratocystidaceae are the respective symbionts of three independent lineages of ambrosia beetles with large, complex mycangia. Fungal Biology, 119(11), 1075–1092.

38. Moran, N. A., McCutcheon, J. P., & Nakabachi, A. (2008). Genomics and evolution of heritable bacterial symbionts. Annual review of genetics, 42(1), 165–190.

39. Moran, N. A., & Sloan, D. B. (2015). The hologenome concept: helpful or hollow? PLoS Biology, 13(12), e1002311.

40. Mueller, U. G., Schultz, T. R., Currie, C. R., Adams, R. M., & Malloch, D. (2001). The origin of the attine ant–fungus mutualism. Quarterly Review of Biology, 76, 169–197.

41. Nowak, M. A. (2006). Evolutionary dynamics: exploring the equations of life. Harvard University Press.

42. Nowak, M. A., & Sigmund, K. (2004). Evolutionary dynamics of biological games. Science, 303, 793–799.

43. O’Donnell, K., Libeskind-Hadas, R., Hulcr, J., et al. (2015). Repeated host shifts in the Fusarium–Euwallacea mutualism. Fungal Genetics and Biology, 82, 277–290.

44. Otto, S. P., & Whitlock, M. C. (1997). The probability of fixation in populations of changing size. Genetics, 146, 723–733.

45. Rocha, E. P. (2018). Neutral theory, microbial practice: challenges in bacterial population genetics. Molecular Biology and Evolution, 35(6), 1338–1347.

46. Six, D.L. (2012). Ecological and evolutionary determinants of bark beetle–fungus symbioses. Insects, 3, 339–366.

47. Taylor, P. D., & Jonker, L. B. (1978). Evolutionary stable strategies and game dynamics. Mathematical Biosciences, 40, 145–156.

48. Vega, F. E., & Blackwell, M. (Eds.). (2005). Insect-fungal associations: ecology and evolution. Oxford University Press.

49. Volterra, V. (1927). Fluctuations in the abundance of a species considered mathematically. Nature, 119, 12–13.

50. Wernegreen, J.J. (2015). Endosymbiont evolution: predictions from theory and surprises from genomes. Annals of the New York Academy of Sciences, 1360, 16–35.

